# Neutralization of Omicron sublineages and Deltacron SARS-CoV-2 by 3 doses of BNT162b2 vaccine or BA.1 infection

**DOI:** 10.1101/2022.06.05.494889

**Authors:** Chaitanya Kurhade, Jing Zou, Hongjie Xia, Mingru Liu, Qi Yang, Mark Cutler, David Cooper, Alexander Muik, Ugur Sahin, Kathrin U. Jansen, Ping Ren, Xuping Xie, Kena A. Swanson, Pei-Yong Shi

## Abstract

Distinct SARS-CoV-2 Omicron sublineages have evolved showing increased fitness and immune evasion than the original Omicron variant BA.1. Here we report the neutralization activity of sera from BNT162b2 vaccinated individuals or unimmunized Omicron BA.1-infected individuals against Omicron sublineages and “Deltacron” variant (XD). BNT162b2 post-dose 3 immune sera neutralized USA-WA1/2020, Omicron BA.1-, BA.2-, BA.2.12.1-, BA.3-, BA.4/5-, and XD-spike SARS-CoV-2s with geometric mean titers (GMTs) of 1335, 393, 298, 315, 216, 103, and 301, respectively; thus, BA.4/5 SARS-CoV-2 spike variant showed the highest propensity to evade vaccine neutralization compared to the original Omicron variants BA.1. BA.1-convalescent sera neutralized USA-WA1/2020, BA.1-, BA.2-, BA.2.12.1-, BA.3-, BA.4/5-, and Deltacron-spike SARS-CoV-2s with GMTs of 15, 430, 110, 109, 102, 25, and 284, respectively. The low neutralization titers of vaccinated sera or convalescent sera from BA. 1 infected individuals against the emerging and rapidly spreading Omicron BA.4/5 variants provide important results for consideration in the selection of an updated vaccine in the current Omicron wave.

## TO THE EDITOR

The severe acute respiratory syndrome coronavirus 2 (SARS-CoV-2) Omicron variant has emerged as the fifth variant of concern (VOC) after Alpha, Beta, Gamma, and Delta. Omicron exhibited the greatest evasion from vaccine- and infection-elicited neutralizing antibodies among all the VOCs.^1^ Since the emergence of Omicron in late November 2021, distinct sublineages have evolved around the world: BA.1 was responsible for the initial surge and has now been replaced by BA.2 and BA.2.12.1 in the USA (https://covid.cdc.gov/covid-data-tracker/#variant-proportions); BA.4 and BA.5 have become prevalent in South Africa and, more recently, are becoming prevalent in some areas in Europe and are increasing in the USA; BA.3 currently remains at low frequency. The co-circulation of Omicron and Delta has also led to a hybrid variant “Deltacron”. Since these newly emerged Omicron sublineages and Deltacron have distinct mutations in the spike glycoproteins (**Fig. S1A**), it is important to examine their susceptibility to vaccine- or infection-elicited antibody neutralization.

To assess the neutralization titers of sera from BNT162b2-immunized or Omicron BA.1-infected individuals against different variants, we engineered the complete spike gene from Omicron sublineages BA.1 (GISAID EPI_ISL_6640916), BA.2 (GISAID EPI_ISL_6795834.2), BA.2.12.1 (GISAID EPI_ISL_12115772), BA.3 (GISAID EPI_ISL_7605591), BA.4/5 (BA.4: GISAID EPI_ISL_11542270; BA.5: GISAID EPI_ISL_11542604; BA.4 and BA.5 have the identical spike sequence), and Deltacron XD strain (GISAID EPI_ISL_10819657 with the first 158 amino acids from Delta spike and the rest from Omicron BA.1 spike) into the mNeonGreen (mNG) reporter USA-WA1/2020 strain, a SARS-CoV-2 strain isolated in January 2020 (**Fig. S1A**). The resulting BA.1-, BA.2-, BA.2.12.1-, BA.3-, BA.4/5-, and XD-spike mNG SARS-CoV-2s produced infectious titers in Vero E6 cells of >10^6^ plaque-forming units per milliliter (PFU/ml), similar to the wild-type USA-WA1/2020 mNG virus. The recombinant viruses developed decreasing sizes of fluorescent foci in the order of USA-WA1/2020 > BA.4/5-spike > BA.2-spike ≈ BA.2.12.1-spike ≈ BA.3-spike > XD-spike ≈ BA.1-spike mNG SARS-CoV-2 (**Fig. S1B**). All recombinant viruses were sequenced to ensure no undesired mutations. Only the sequenced passage 1 viruses were used for the neutralization assays.

Using the recombinant SARS-CoV-2s, we determined the 50% fluorescent focus-reduction neutralization titers (FFRNT_50_) for two panels of human sera. The first serum panel (n=22) was collected from individuals at 1-month post-dose 3 (PD3) of BNT162b2 vaccine. We chose the PD3 sera because (i) many people have already received 3 doses of BNT162b2 and (ii) two doses of BNT162b2 did not elicit robust neutralization against Omicron BA.1.^2^ The second serum panel (n=20) was collected from unvaccinated COVID-19 patients who had been infected by BA.1 during the initial Omicron emergence in the USA.^3^ The genotype of the infecting virus was verified for each patient by Sanger sequencing of nasal swabs. **Tables S1** and **S2** summarize the data from the PD3 BNT162b2-vaccinated- and BA.1-infected individuals, respectively.

BNT162b2 PD3 immune sera neutralized USA-WA1/2020, BA.1-, BA.2-, BA.2.12.1-, BA.3-, BA.4/5-, and XD-spike mNG SARS-CoV-2s with geometric mean titers (GMTs) of 1335, 393, 298, 315, 216, 103, and 301, respectively (**Fig. 1A** and **Table S1**). All PD3 sera neutralized all variant-spike SARS-CoV-2s with neutralization titers of >20.

**Figure 1.**
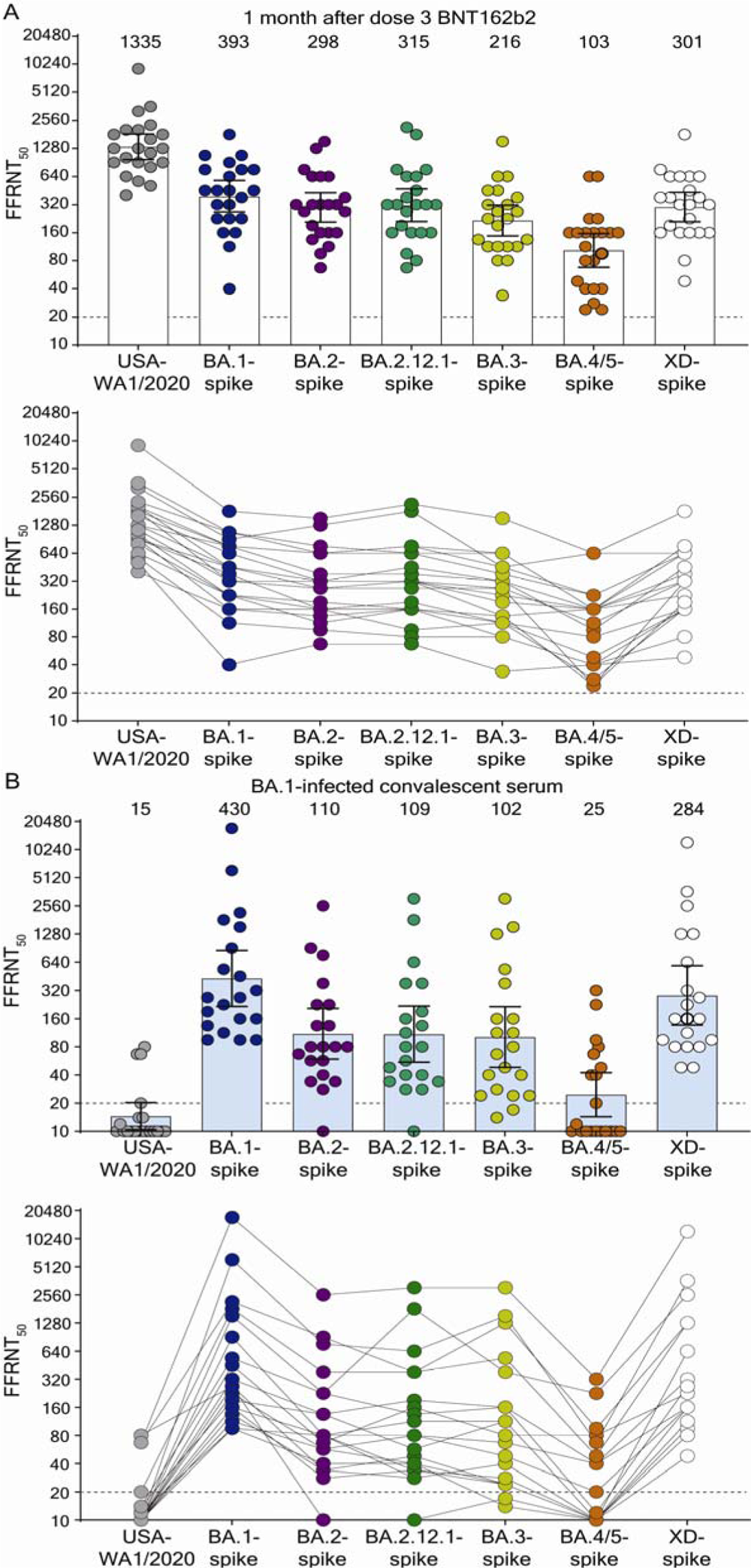
Neutralization by sera collected at 1 month post dose 3 BNT162b2 vaccine (A) and by sera collected from unvaccinated individuals who contracted Omicron BA.1 SARS-CoV-2 (B). Scatterplot of neutralization titers against USA-WA1/2020, Omicron sublineage BA.1-, BA.2-, BA.2.12.1-, BA.3-, BA.4/5-, and Deltacron XD-spike mNG SARS-CoV-2s. Both BNT162b2-vaccinated sera (n=22) and BA.1-infected convalescent sera (n=20) were tested for their FFRNT_50_s against the variant-spike mNG SARS-CoV-2s. The variant-spike mNG SARS-CoV-2s were produced by engineering the complete variant spike genes into the mNG USA-WA1/2020. Each data point represents the geometric mean FFRNT_50_ (GMT) obtained with a serum specimen against the indicated virus. **Tables S1** and **S2** summarize the serum information and FFRNT_50_s for (**A**) and (**B**), respectively. The neutralization titers for individual variant-spike mNG SARS-CoV-2s were determined in two or three independent experiments, each with duplicate assays; the GMTs are presented. The bar heights and the numbers above indicate GMTs. The whiskers indicate 95% confidence intervals. The dotted lines indicate the limit of detection of FFRNT_50_. Statistical analysis was performed with the use of the Wilcoxon matched-pairs signed-rank test. For the BNT162b2-vaccinated sera in (**A**), the *P* values of the GMT differences between USA-WA1/2020 and any variant-spike SARS-CoV-2 are all < 0.0001. For the BA.1-convelescent sera in (**B**), the *P* value of GMT difference between BA.1- and XD-spike viruses is 0.0021; the *P* values of the GMT differences between BA.1- and any other variant-spike viruses (including USA-WA1/2020) are all <0.0001. For both serum panels in (**A**) and (**B**), FFRNT_50_ values with connected lines are presented for individual sera.

However, neutralizing GMTs against BA.1-, BA.2-, BA.2.12.1-, BA.3-, BA.4/5-, and XD-spike SARS-CoV-2s were 3.4-, 4.5-, 4.2-, 6.2-, 13.0-, and 4.4-fold lower than that against USA-WA1/2020 GMT, respectively (**Fig. 1A**). The results support two conclusions. First, BA.1, BA.2, BA.2.12.1, and XD spike variants have an increased but similar propensity to evade neutralizing antibodies induced by 3 doses of BNT162b2 and likely other COVID-19 vaccines. This result suggests that the sequential increase in the prevalence of BA.2.12.1 > BA.2 > BA.1 over the past 6 months was likely not driven by the difference in spike-mediated evasion of neutralization in vaccinated people, but by other factors, such as differences in viral transmission, or other mechanisms of immune evasion. Second, BA.4/5 variants are much less efficiently neutralized by BNT162b2 PD3 immune sera than BA.1, BA.2, BA.2.12.1, BA.3 and, XD spikes. The BA.3 sublineage may be attenuated in viral fitness as it has remained at very low prevalence; however, BA.4 and BA.5 sublineages, with the additional F486V and reversion of Q493 (compared to BA.2.12.1) that appear to improve infectivity, may potentially replace other Omicron sublineages in circulation. These findings suggest that close monitoring of the prevalence of sublineages BA.4 and BA.5 through epidemiological surveillance is critical as rapid increases in these sublineages with higher levels of immune escape could lead to new waves of infections.

Sera from BA.1-infected individuals neutralized USA-WA1/2020, BA.1-, BA.2-, BA.2.12.1-, BA.3-, BA.4/5-, and XD-spike viruses with GMTs of 15, 430, 110, 109, 102, 25, and 284, respectively (**Fig. 1B** and **Table S2**). Thus, the neutralizing GMTs against heterologous USA-WA1/2020-, BA.2-, BA.2.12.1-, BA.3-, BA.4/5-, and XD-spike SARS-CoV-2s were 28.7-, 3.9-, 3.9-, 4.2-, 17.2-, and 1.5-fold lower than that against homologous BA.1-spike virus, respectively. All sera neutralized BA.1-spike virus with titers of ≥80. Importantly, however, 13 (56%), 1 (5%), 1 (5%), and 10 (50%) out of 20 sera did not neutralize USA-WA1/2020, BA.2-, BA.2.12.1-, and BA.4/5-spike SARS-CoV-2s, respectively (**Fig. 1B**). These results suggest that sera from BA.1-infected individuals do not neutralize all Omicron sublineages and Deltacron XD with similar efficiencies, and importantly are not efficient at neutralizing BA.4/5. The addtitional mutation F486V in the BA.4/5 spike may account for the further evasion of neutralization (**Fig. S1A**).

The current study has confirmed and expanded our previous results on Omicron sublineage neutralization.^3^ Our results have potential implications for the strategy of updating the current COVID-19 vaccines in which, even within the Omicron lineages, substantial differences exist in the potential of vaccinated or BA.1-infected individuals to neutralize Omicron sublineages. The data suggest that Omicron BA.4 and BA.5 sublineages are adapted to evade neutralization in both prototype vaccinated and Omicron BA.1-infected individuals. The antigenic distinctions of spike glycoproteins among Omicron sublineages, Deltacron XD, and USA-WA1/2020 as well as prevalence of Omicron sublineages must be carefully considered when deciding to update the vaccine to new variants.

Our study has some limitations. This study lacks analyses of T cells and non-neutralizing antibodies that can mediate Fc-mediated effector functions. Both of these immune arms, in conjunction with neutralizing antibodies, protect patients from severe disease. After vaccination or infection, the majority of the T cell epitopes are preserved in Omicron spikes.^4^ Caution should also be taken when comparing the differences in variant neutralization between BNT162b2-vaccinated (PD3 sera) and BA.1-infected individuals. This is because (i) a third dose booster vaccination increases both the magnitude and breadth of neutralization;^5^ (ii) BA.1-infected sera used in this study (collected on days 8 to 62 after a positive RT-PCR test) were heterogeneous, with some sera collected at an acute plasma blast stage and other sera at a convalescent IgG dominant phase; and (iii) the relatively small sample size. Regardless, our results consistently showed the most significant reduction in neutralization against BA.4/5 spike, underscoring the importance of closely monitoring the prevalence of sublineages BA.4 and BA.5 globally and taking the data into account when making decisions on updating current COVID-19 vaccines.

## Methods

### Human Sera

Two human serum panels were used in the study. The first panel of serum samples was collected from BNT162b2 vaccinees participating in the phase 1 portion of the ongoing phase 1/2/3 clinical trial (ClinicalTrials.gov identifier: NCT04368728). The protocol and informed consent were approved by institutional review boards for each of the investigational centers participating in the study. The study was conducted in compliance with all International Council for Harmonisation Good Clinical Practice guidelines and the ethical principles of the Declaration of Helsinki. The primary outcomes for phase 1 were reported previously.^1,6,7^ BNT162b2-vaccinated sera (n=22) were collected 1 month after dose 3 and used for neutralization test. **Table S1** summarizes the patient information which was previously reported.^5,8^

The second serum panel (n=20) was collected from COVID-19 patients who were not vaccinated but infected by BA.1 sublineage during the initial Omicron emergence at the University of Texas Medical Branch (UTMB).^3^ The research protocol regarding the use of human sera was reviewed and approved by the UTMB Institutional Review Board (IRB number 20-0070). The de-identified human sera were heat-inactivated at 56°C for 30 min before the neutralization test. The genotype of infecting virus was verified by Sanger sequencing. **Table S2** summarizes the serum information which was recently reported.^3^

### Recombinant Omicron sublineage- or Deltacron XD-spike mNG SARS-CoV-2s

Recombinant Omicron sublineage BA.1-, BA.2-, BA.2.12.1-, BA.3-, BA.4/5-, and Deltacron XD-spike mNG SARS-CoV-2s were constructed by engineering the complete spike gene from the indicated variants into an infectious cDNA clone of mNG USA-WA1/2020^9^ (**Fig. S1A**). All amino acid substitutions, deletions, and insertions in the variant spike glycoproteins were introduced into the infectious cDNA clone of mNG USA-WA1/2020 using PCR-based mutagenesis as previously described.^10^ The BA.1-, BA.2-, BA.2.12.1-, BA.3-, BA4/5-, and XD-spike sequences were based on GISAID EPI_ISL_6640916, EPI_ISL_6795834.2, EPI_ISL_12115772, EPI_ISL_7605591, EPI_ISL_11542270 (BA.4 spike is identical to BA.5 spike EPI_ISL_11542604), and EPI_ISL_10819657, respectively. **Figure S1A** depicts the spike mutations from different Omicron sublineages and Deltacron XD variant. The full-length cDNA of viral genome bearing the variant spike was assembled via *in vitro* ligation and used as a template for *in vitro* transcription. The full-length viral RNA was then electroporated into Vero E6-TMPRSS2 cells. On day 3-4 post electroporation, the original P0 virus was harvested from the electroporated cells and propagated for another round on Vero E6 cells to produce the P1 virus. The infectious titer of the P1 virus was quantified by fluorescent focus assay on Vero E6 cells (**Fig. S1B**) and sequenced for the complete spike gene to ensure no undesired mutations. The P1 virus was used for the neutralization test. The protocols for the mutagenesis of mNG SARS-CoV-2 and virus production were reported previously.^11^

### Fluorescent focus reduction neutralization test (FFRNT)

FFRNT was performed to measure the neutralization titers of sera against USA-WA1/2020, BA.1-, BA.2-, BA.2.12.1-, BA.3-, BA4/5-, and XD-spike mNG SARS-CoV-2s. The FFRNT protocol was reported previously.^12^ Vero E6 cells were seeded onto 96-well plates with 2.5×10^4^ cells per well (Greiner Bio-one™) and incubated overnight. On the next day, each serum was 2-fold serially diluted in culture medium and mixed with 100-150 FFUs of mNG SARS-CoV-2. The final serum dilution ranged from 1:20 to 1:20,480. After incubation at 37°C for 1 h, the serum-virus mixtures were loaded onto the pre-seeded Vero E6 cell monolayer in 96-well plates. After 1 h infection, the inoculum was removed and 100 μl of overlay medium containing 0.8% methylcellulose was added to each well. After incubating the plates at 37°C for 16 h, raw images of mNG foci were acquired using Cytation™ 7 (BioTek) armed with 2.5× FL Zeiss objective with a wide-field of view and processed using the software settings (GFP [469,525] threshold 4000, object selection size 50-1000 μm). The fluorescent mNG foci were counted in each well and normalized to the non-serum-treated controls to calculate the relative infectivities. The FFRNT_50_ value was defined as the minimal serum dilution to suppress >50% of fluorescent foci. The neutralization titer of each serum was determined in duplicate assays, and the geometric mean was taken. **Tables S1** and **S2** summarize the FFRNT50 results.

## Acknowledgments

We thank the Pfizer-BioNTech clinical trial C4591001 and NCT04368728 participants, from whom the post-immunization human sera were obtained. We thank many colleagues at Pfizer and BioNTech who developed and produced the BNT162b2 vaccine. The collection and testing of BA.1-infected human sera at UTMB were supported by NIH grant HHSN272201600013C and an award from the Sealy & Smith Foundation.

## Author contributions

K.U.J., P.R., X.X., K.A.S., and P.-Y.S. conceived the study. C.K., J.Z., H.X., M.L., Q.Y., M.C., D.C., P.R., and X.X. performed the experiments. C.K., J.Z., M.C., D.C., A.M., K.U.J., P.R., X.X., K.A.S., and P.-Y.S. analyzed the results. C.K., J.Z., P.R., X.X., K.A.S., and P. -Y.S. wrote the manuscript. D.C., K.U.J., P.R., X.X., K.A.S., and P.-Y.S. supervised the project.

