## Supplemental figure1 and table S1-2 for "Neutralization of Omicron sublineages and Deltacron SARS-CoV-2 by 3 doses of BNT162b2 vaccine or BA.1 infection"

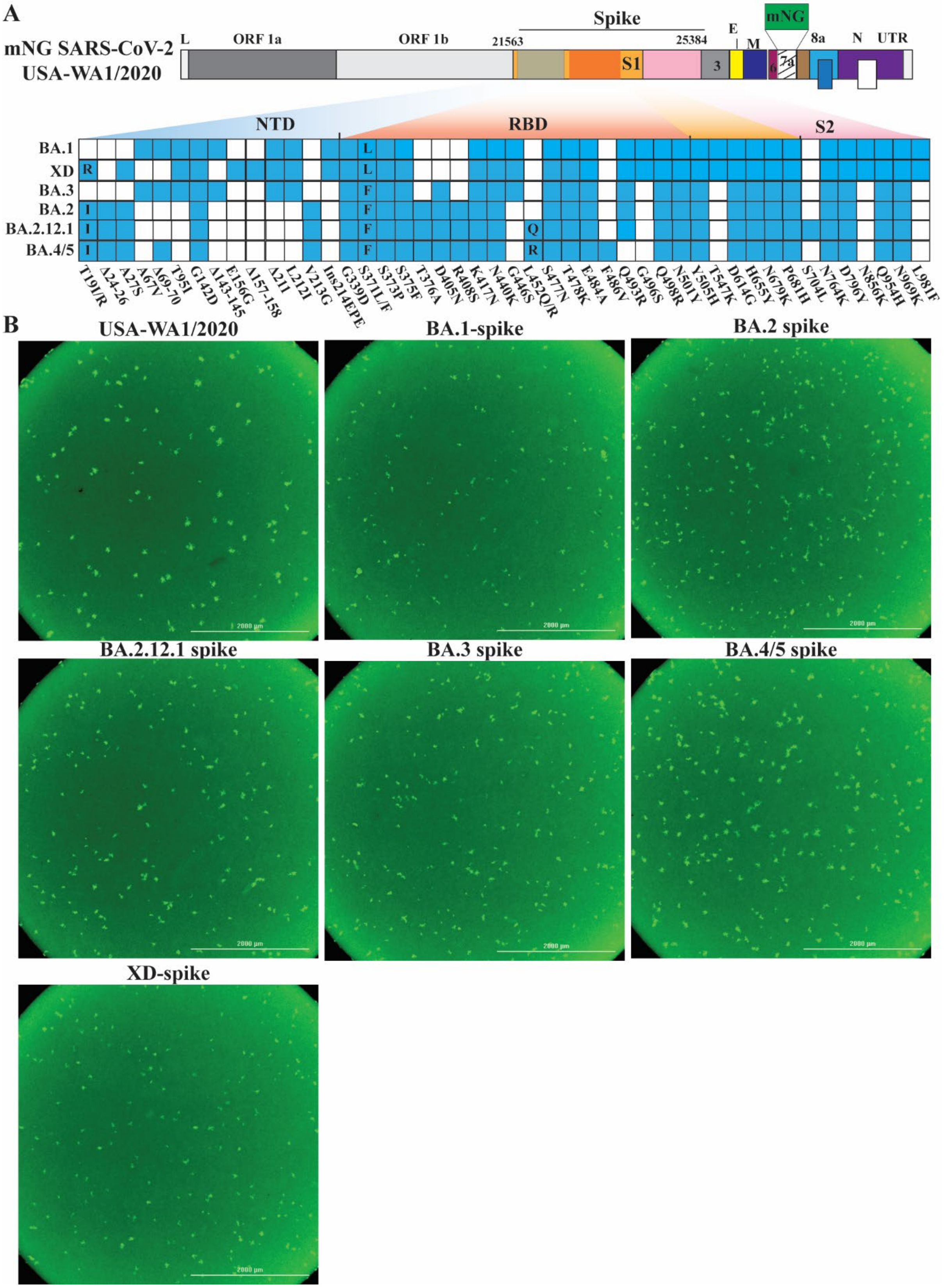
A
mNG SARS-CoV-2
USA-WA1/2020
L
ORF 1a
ORF 1b
21563
Spike
25384
S1
3
E
M
mNG
6
7a
8a
N
UTR
NTD
RBD
S2
BA.1
XD
BA.3
BA.2
BA.2.12.1
BA.4/5
T19I/R
Δ24-26
A27S
A67V
Δ69-70
T95I
G142D
Δ143-145
E156G
Δ157-158
Δ211
L212I
V213G
Ins214EPE
G339D
S371L/F
S373P
S375F
T376A
D405N
R408S
K417N
N440K
G446S
L452Q/R
S477N
T478K
E484A
F486V
Q493R
G496S
Q498R
N501Y
Y505H
T547K
D614G
H655Y
N679K
P681H
S704L
N764K
D796Y
N856K
Q954H
N969K
L981F
B
USA-WA1/2020
BA.1-spike
BA.2 spike
BA.2.12.1 spike
BA.3 spike
BA.4/5 spike
XD-spike
2000 µm

**Figure S1. Construction of Omicron sublineage BA.1-, BA.2-, BA.2.12.1-, BA.3-, BA.4/5-, and Deltacron XD-spike mNG SARS-CoV-2s.** (**A**) Omicron sublineage- and Deltacron XD-spike mNG SARS-CoV-2s. mNG USA-WA1/2020 SARS-CoV-2 was used to engineer Omicron sublineage- and Deltacron XD-spike SARS-CoV-2s. The mNG reporter gene was engineered at the open-reading-frame-7 (ORF7) of the USA-WA1/2020 genome.[1] Amino acid mutations, deletions (Δ), and insertions (Ins) are indicated for variant spikes in reference to the USA-WA1/2020 spike. L: leader sequence; ORF: open reading frame; NTD: N-terminal domain of S1; RBD: receptor binding domain of S1; S: spike glycoprotein; S1: N-terminal furin cleavage fragment of S; S2: C-terminal furin cleavage fragment of S; E: envelope protein; M: membrane protein; N: nucleoprotein; UTR: untranslated region. (**B**) Fluorescent foci formed by USA-WA1/2020, BA.1-, BA.2-, BA.2.12.1-, BA.3-, BA.4/5- and XD-spike mNG SARS-CoV-2s. The images were taken at 16 h after Vero E6 cells were infected with the indicated viruses in 96-well plates. Representative images are presented for each virus.

### Table S1. $FFRNT_{50}$ values of BNT162b2-vaccinated sera

| Subject [#]ID | Age (Years) | Sex (F/M) | *$FFRNT_{50}$ | | | | | | | | | | | | | | | | | | | | |
|---|---|---|---|---|---|---|---|---|---|---|---|---|---|---|---|---|---|---|---|---|---|---|---|
| | | | USA-WA1/2022 | | | | BA.1-spike | | | BA.2-spike | | | BA.2.12.1-spike | | | BA.3-spike | | | BA.4/5-spike | | | XD-spike | |
| | | | [&]Exp1 | [&]Exp2 | Exp3 | GMT | [&]Exp1 | Exp2 | GMT | [&]Exp1 | Exp2 | GMT | Exp1 | Exp2 | GMT | [&]Exp1 | Exp2 | GMT | Exp1 | Exp2 | GMT | Exp1 | Exp2 | GMT |
| 1 | 26 | F | 640 | 905 | 1280 | 905 | 320 | 113 | 453 | 320 | 320 | 320 | 320 | 320 | 320 | 226 | 226 | 226 | 160 | 160 | 160 | 320 | 453 | 381 |
| 2 | 28 | M | 1280 | 1810 | 2560 | 1810 | 320 | 640 | 320 | 320 | 320 | 320 | 320 | 320 | 320 | 113 | 160 | 135 | 160 | 160 | 160 | 160 | 320 | 226 |
| 3 | 35 | F | 1280 | 1280 | 1280 | 1280 | 640 | 40 | 761 | 453 | 57 | 381 | 320 | 453 | 381 | 320 | 320 | 320 | 113 | 80 | 95 | 320 | 640 | 453 |
| 4 | 35 | F | 1280 | 1280 | 1280 | 1280 | 320 | 640 | 453 | 226 | 320 | 269 | 226 | 320 | 269 | 226 | 320 | 269 | 20 | 28 | 24 | 320 | 320 | 320 |
| 5 | 38 | F | 1280 | 1810 | 2560 | 1810 | 640 | 1280 | 761 | 640 | 905 | 640 | 640 | 640 | 640 | 320 | 453 | 381 | 226 | 226 | 226 | 640 | 640 | 640 |
| 6 | 38 | F | 320 | 320 | 640 | 403 | 160 | 226 | 160 | 160 | 160 | 160 | 80 | 113 | 95 | 80 | 160 | 113 | 28 | 28 | 28 | 160 | 160 | 160 |
| 7 | 44 | F | 640 | 453 | 640 | 570 | 113 | 320 | 113 | 113 | 113 | 95 | 80 | 80 | 80 | 80 | 80 | 80 | 40 | 40 | 40 | 80 | 80 | 80 |
| 8 | 44 | F | 2560 | 1280 | 2560 | 2032 | 640 | 640 | 761 | 640 | 320 | 640 | 640 | 905 | 761 | 320 | 640 | 453 | 226 | 226 | 226 | 640 | 640 | 640 |
| 9 | 52 | F | 2560 | 1280 | 2560 | 2032 | 320 | 905 | 453 | 320 | 640 | 320 | 453 | 453 | 453 | 320 | 320 | 320 | 160 | 160 | 160 | 453 | 320 | 381 |
| 10 | 54 | M | 1280 | 1280 | 1280 | 1280 | 453 | 160 | 453 | 320 | 160 | 320 | 320 | 320 | 320 | 226 | 320 | 269 | 80 | 160 | 113 | 320 | 320 | 320 |
| 11 | 65 | M | 1280 | 640 | 1280 | 1016 | 226 | 453 | 226 | 226 | 160 | 190 | 160 | 160 | 160 | 113 | 113 | 113 | 80 | 80 | 80 | 160 | 160 | 160 |
| 12 | 65 | M | 5120 | 2560 | 2560 | 3225 | 640 | 905 | 905 | 1280 | 320 | 1280 | 1810 | 1810 | 1810 | 320 | 640 | 453 | 640 | 640 | 640 | 640 | 640 | 640 |
| 13 | 66 | F | 640 | 640 | 1280 | 806 | 160 | 1280 | 226 | 160 | 640 | 160 | 226 | 160 | 190 | 80 | 160 | 113 | 80 | 80 | 80 | 160 | 160 | 160 |
| 14 | 67 | M | 640 | 905 | 1280 | 905 | 320 | 320 | 381 | 160 | 320 | 160 | 160 | 160 | 160 | 113 | 160 | 135 | 28 | 20 | 24 | 160 | 226 | 190 |
| 15 | 68 | M | 640 | 905 | 1280 | 905 | 160 | 2560 | 160 | 113 | 1280 | 135 | 160 | 160 | 160 | 113 | 113 | 113 | 40 | 57 | 48 | 160 | 160 | 160 |
| 16 | 68 | F | 5120 | 2560 | 3620 | 3620 | 905 | 160 | 1076 | 640 | 160 | 640 | 640 | 905 | 761 | 640 | 640 | 640 | 160 | 160 | 160 | 640 | 905 | 761 |
| 17 | 68 | F | 640 | 640 | 640 | 640 | 160 | 320 | 226 | 113 | 160 | 113 | 160 | 160 | 160 | 80 | 80 | 80 | 40 | 40 | 40 | 160 | 160 | 160 |
| 18 | 69 | F | 1280 | 905 | 1280 | 1140 | 320 | 1280 | 320 | 226 | 1280 | 269 | 320 | 320 | 320 | 160 | 226 | 190 | 160 | 160 | 160 | 320 | 320 | 320 |
| 19 | 69 | F | 2560 | 1810 | 2560 | 2281 | 905 | 320 | 1076 | 640 | 320 | 761 | 640 | 640 | 640 | 640 | 640 | 640 | 160 | 160 | 160 | 640 | 640 | 640 |
| 20 | 70 | M | 10240 | 10240 | 7241 | 9123 | 1280 | 640 | 1810 | 1810 | 320 | 1522 | 2560 | 1810 | 2153 | 1810 | 1280 | 1522 | 640 | 640 | 640 | 1810 | 1810 | 1810 |
| 21 | 73 | M | 320 | 453 | 640 | 508 | 40 | 453 | 40 | 80 | 320 | 67 | 57 | 80 | 67 | 28 | 40 | 34 | 40 | 40 | 40 | 40 | 57 | 48 |
| 22 | 74 | F | 1280 | 1280 | 2560 | 1613 | 640 | 905 | 640 | 320 | 640 | 320 | 320 | 320 | 320 | 160 | 320 | 226 | 160 | 160 | 160 | 320 | 453 | 381 |
| [†]GMT | -- | -- | 1280 | 1146 | 1596 | 1335 | 336 | 460 | 393 | 300 | 315 | 298 | 305 | 325 | 315 | 190 | 245 | 216 | 101 | 105 | 103 | 282 | 320 | 301 |
| [^]95% CI | -- | -- | 866-1893 | 827-1589 | 1208-2109 | 972-1834 | 234-482 | 301-702 | 266-578 | 210-431 | 223-446 | 208-428 | 202-462 | 219-483 | 211-471 | 127-284 | 169-356 | 148-316 | 66-155 | 69-159 | 68-156 | 195-409 | 223-461 | 209-432 |

*Individual $FFRNT_{50}$ value is the geometric mean of duplicate FFRNT results.
[#]The sera were collected at 1 month post dose 3 BNT162b2 vaccine, as recently reported.[2]
[†]Geometric mean neutralizing titer (GMT).
[^]95% confidence interval (95% CI) for the GMT.
[&]These data were reported previously.[3]

### Table S2. FFRNT$_{50}$ values of Omicron sublineage BA.1-infected sera

| Serum ID | Age (years) | Gender (F/M) | *FFRNT$_{50}$ | | | | | | | | | | | | | | | | | | | | | Serum collection time (days post positive RT-PCR test) |
|---|---|---|---|---|---|---|---|---|---|---|---|---|---|---|---|---|---|---|---|---|---|---|---|---|
| | | | USA-WA1/2020 | | | BA.1-spike | | | BA.2-spike | | | BA.2.12.1-spike | | | BA.3-spike | | | BA.4/5-spike | | | XD-spike | | | |
| | | | [&]Exp1 | Exp2 | GMT | [&]Exp1 | Exp2 | GMT | [&]Exp1 | Exp2 | GMT | Exp1 | Exp2 | GMT | [&]Exp1 | Exp2 | GMT | Exp1 | Exp2 | GMT | Exp1 | Exp2 | GMT | |
| 1 | 21-30 | M | ^10 | 10 | 10 | 80 | 113 | 95 | 160 | 40 | 80 | 40 | 28 | 34 | 20 | 28 | 24 | 10 | 10 | 10 | 80 | 113 | 95 | 26 |
| 2 | 41-50 | F | 10 | 10 | 10 | 80 | 113 | 95 | 57 | 57 | 80 | 40 | 20 | 28 | 40 | 40 | 40 | 10 | 10 | 10 | 80 | 80 | 80 | 33 |
| 3 | 81-90 | M | 10 | 10 | 10 | 113 | 160 | 135 | 80 | 40 | 34 | 57 | 40 | 48 | 28 | 20 | 24 | 10 | 10 | 10 | 80 | 113 | 95 | 21 |
| 4 | 11-20 | M | 14 | 10 | 12 | 113 | 80 | 95 | 160 | 40 | 40 | 40 | 40 | 40 | 14 | 14 | 14 | 10 | 10 | 10 | 40 | 57 | 48 | 16 |
| 5 | 31-40 | F | 10 | 10 | 10 | 160 | 320 | 226 | 113 | 80 | 57 | 80 | 80 | 80 | 40 | 57 | 48 | 20 | 20 | 20 | 160 | 160 | 160 | 16 |
| 6 | 21-30 | F | 10 | 10 | 10 | 160 | 160 | 160 | 28 | 40 | 28 | 40 | 28 | 34 | 20 | 40 | 28 | 10 | 10 | 10 | 80 | 80 | 80 | 28 |
| 7 | 1-10 | F | 10 | 10 | 10 | 160 | 160 | 160 | 10 | 113 | 80 | 80 | 80 | 80 | 57 | 80 | 67 | 40 | 40 | 40 | 160 | 160 | 160 | 43 |
| 8 | 61-70 | M | 10 | 10 | 10 | 160 | 80 | 113 | 226 | 40 | 57 | 40 | 40 | 40 | 20 | 28 | 24 | 10 | 10 | 10 | 80 | 80 | 80 | 40 |
| 9 | 81-90 | M | 20 | 10 | 14 | 226 | 160 | 190 | 1280 | 113 | 135 | 57 | 57 | 57 | 226 | 57 | 113 | 10 | 10 | 10 | 113 | 113 | 113 | 40 |
| 10 | 1-10 | F | 10 | 10 | 10 | 320 | 226 | 269 | 160 | 40 | 34 | 28 | 28 | 28 | 40 | 40 | 40 | 10 | 10 | 10 | 160 | 160 | 160 | 26 |
| 11 | 1-10 | F | 10 | 10 | 10 | 320 | 320 | 320 | 28 | 160 | 135 | 160 | 226 | 190 | 160 | 160 | 160 | 40 | 40 | 40 | 320 | 320 | 320 | 56 |
| 12 | 51-60 | F | 10 | 10 | 10 | 453 | 453 | 453 | 20 | 160 | 67 | 160 | 160 | 160 | 80 | 80 | 80 | 80 | 80 | 80 | 226 | 226 | 226 | 35 |
| 13 | 71-80 | M | 160 | 40 | 80 | 453 | 160 | 269 | 2560 | 10 | 10 | 10 | 10 | 10 | 20 | 14 | 17 | 10 | 10 | 10 | 57 | 40 | 48 | 17 |
| 14 | 21-30 | F | 20 | 10 | 14 | 640 | 453 | 538 | 40 | 113 | 226 | 113 | 160 | 135 | 160 | 160 | 160 | 14 | 10 | 12 | 226 | 320 | 269 | 62 |
| 15 | 51-60 | M | 10 | 10 | 10 | 905 | 905 | 905 | 40 | 160 | 80 | 160 | 80 | 113 | 113 | 113 | 113 | 10 | 10 | 10 | 640 | 640 | 640 | 29 |
| 16 | 71-80 | F | 10 | 10 | 10 | 1280 | 1810 | 1522 | 453 | 226 | 226 | 1280 | 2560 | 1810 | 320 | 453 | 381 | 226 | 226 | 226 | 2560 | 2560 | 2560 | 8 |
| 17 | 71-80 | F | 28 | 14 | 20 | 2560 | 1280 | 1810 | 453 | 320 | 381 | 320 | 453 | 381 | 640 | 453 | 538 | 57 | 40 | 48 | 1280 | 1280 | 1280 | 32 |
| 18 | 61-70 | M | 113 | 40 | 67 | 2560 | 1810 | 2153 | 640 | 640 | 905 | 320 | 453 | 381 | 1810 | 905 | 1280 | 80 | 113 | 95 | 1280 | 1280 | 1280 | 15 |
| 19 | 71-80 | F | 10 | 10 | 10 | 5120 | 7241 | 6089 | 113 | 905 | 761 | 640 | 640 | 640 | 1280 | 1810 | 1522 | 80 | 57 | 67 | 2560 | 5120 | 3620 | 16 |
| 20 | 81-90 | M | 40 | 113 | 67 | 14482 | 20480 | 17222 | 28 | 2560 | 2560 | 2560 | 3620 | 3044 | 2560 | 3620 | 3044 | 320 | 320 | 320 | 10240 | 14482 | 12177 | 13 |
| [#]GMT | - | - | 16 | 13 | 15 | 445 | 415 | 430 | 115 | 113 | 110 | 109 | 109 | 109 | 102 | 102 | 102 | 25 | 24 | 25 | 269 | 299 | 284 | 25 |
| [†]95% CI | - | - | 11-24 | 10-18 | 10-20 | 225-881 | 205-842 | 216-855 | 58-228 | 62-208 | 59-206 | 58-207 | 52-230 | 55-218 | 48-218 | 48-214 | 49-215 | 15-43 | 14-42 | 14-42 | 132-549 | 141-632 | 137-588 | 20-32 |

*Individual FFRNT$_{50}$ value is the geometric mean of duplicate FFRNT results.
^FFRNT$_{50}$ of <20 was treated as 10 for plot purpose and statistical analysis.
[#]Geometric mean neutralizing titer (GMT).
[†]95% confidence interval (95% CI) for the GMT.
[&] These data were reported previously.[4]
